# The impossible challenge of estimating non-existent moments of the Chemical Master Equation

**DOI:** 10.1101/2023.02.08.527667

**Authors:** Vincent Wagner, Nicole Radde

**Affiliations:** Institute for Systems Theory and Automatic Control, University of Stuttgart, Pfaffenwaldring 9, 70569 Stuttgart, Germany; Stuttgart Center for Simulation Science, University of Stuttgart, Pfaffenwaldring 5a, 70569 Stuttgart, Germany

**Keywords:** Chemical Master Equation, Method of Moments, Heavy-tail Distributions, Stochastic Simulation Algorithm, Moment Estimation

## Abstract

**Motivation:** The Chemical Master Equation is a set of linear differential equations that describes the evolution of the probability distribution on all possible configurations of a (bio-)chemical reaction system. Since the number of configurations and therefore the dimension of the CME rapidly increases with the number of molecules, its applicability is restricted to small systems. A widely applied remedy for this challenge are moment-based approaches which consider the evolution of the first few moments of the distribution as summary statistics for the complete distribution. Here, we investigate the performance of two moment-estimation methods for reaction systems whose equilibrium distributions encounter heavy-tailedness and hence do not possess statistical moments.

**Results:** We show that estimation via Stochastic Simulation Algorithm trajectories lose consistency over time and estimated moment values span a wide range of values even for large sample sizes. In comparison, the Method of Moments returns smooth moment estimates but is not able to indicate the nonexistence of the allegedly predicted moments. We furthermore analyze the negative effect of a CME solution’s heavy-tailedness on SSA run times and explain inherent difficulties.

While moment estimation techniques are a commonly applied tool in the simulation of (bio-)chemical reaction networks, we conclude that they should be used with care, as neither the system definition nor the moment estimation techniques themselves reliably indicate the potential heavy-tailedness of the CME’s solution.

## Introduction

Randomness and uncertainty govern nearly every aspect of our life. This especially also holds for the evolution of (bio-)Chemical Reaction Networks (CRN). In this and many other contexts, a very popular coping strategy is based on the formulation of expectations. Generally, an expectation is a deterministic way of describing a stochastic process by calculating the average of several of its outcomes. Mathematically, the notion of expectation manifests in the form of statistical moments. These quantities as well as their estimation are well-understood. Furthermore, their use in describing stochastic processes is provably founded by the central limit theorem: For processes of finite expectation and variance, the theorem states that the average over several independent realizations is normally distributed with a shrinking variance for a growing number of averaged samples. Consequentially, for sufficiently large sample sizes, one is able to describe many expectation estimators by only the two first statistical moments holistically. While this result is remarkable in every aspect, the central limit theorem is still limited to processes, for which single realizations do not constitute a significant portion of the overall observed behavior. Mandelbrot and Taleb (1) assess, that there indeed exist many applications where the central limit theorem must not be used in order to estimate expectation values of stochastic processes and corresponding risks meaningfully. As an example, they state that 63% of financial returns of half a century can be associated with only 10 trading days. While many authors are aware of non-negligible stochastic effects and use this to their advantage to adequately model for instance bursty gene expression (2–4), expectations remain the instrument of choice to describe stochastic processes in various scientific fields. The Chemical Master Equation (CME) is a standard approach to describe the time evolution of CRNs (see e.g. Higham (5)). Its solution is a time-dependent probability distribution over all configurations of the system. Statistical moments of this distribution can be estimated either via Monte Carlo integration over sample paths from the underlying process, which can be generated via the Stochastic Simulation Algorithm (SSA, Gillespie (6, 7)), or by the Method of Moments (MoM, (8)), a differential equation system for the moments that can be solved via numerical integration. On this basis, different kinds of hybrid methods can be created. Some combine SSA sampling and the MoM (9, 10) depending on the size of the simulated chemical populations. Others complement the estimation of statistical moments with approximations of the distribution’s tail (11). We will nevertheless focus on the two fundamental approaches since the hybrid approaches pose additional conditions to the CRN and are therefore not necessarily better suited to answer a scientific question at hand. In this work, SSA sampling and the MoM are applied to a CRN that converges to a heavy-tail distribution and therefore shows wildly stochastic behavior, the most unpredictable type of stochasticity according to Mandelbrot and Taleb (1). The results are compared to results originating from a very well-behaved network. We demonstrate that expectation estimation techniques for CRNs are not capable of indicating wild stochasticity but instead return meaningless results, thereby deceiving inattentive users.

## Systems and Methods

### Expectation-based approximation techniques for the CME solution

We consider a CRN with *N* molecule types that participate in *J* reactions. The configuration space of such a system is a set of system configurations described by a vector 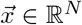 whose entries 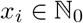 correspond to the number of molecules of species *i*. A reaction *j* is assigned a configuration change vector 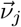, which comprises the changes in molecule numbers if reaction *j* fires once, and a propensity function 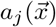. The product 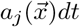 indicates how likely reaction *j* fires in an infinitesimal time instance [*t, t* + *dt*). This defines the CME,

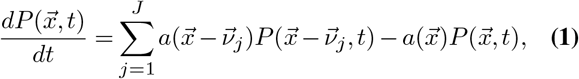

whose solution is a time dependent probability distribution 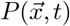 on the space of system configurations. The propensity 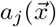 can be chosen by using the law of mass action, according to which the probability of a reaction is proportional to the number of possible educt combinations. The CME describes a time-continuous Markov process on the discrete space of configurations. These processes often converge to an equilibrium distribution, which means that the limit

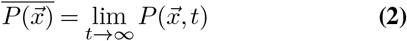

exists independent of the initial condition. This holds for all examples considered in this paper. For clarity and brevity of notation, we restrict our considerations to one-dimensional configuration spaces, i.e. we consider CRNs with one molecular species only. Results can in principle directly be generalized, but the numbers of expectation values and covariances increase linearly and quadratically with the number of molecular species, respectively.

We are particularly interested in the time-dependent first two moments of the probability distribution, the expectation value, and the variance, which are defined via

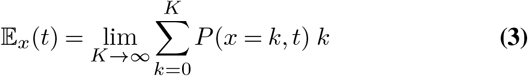

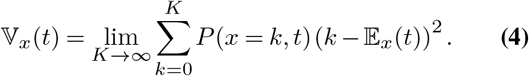

The somewhat lengthy limit formulation stresses that this indeed is a series for which convergence is not trivially given. Throughout this article, we refer to distributions without converging moment series as heavy-tail distributions. Two complementary approaches approximate 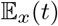 and 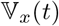. First, moments are estimated via simulating sample trajectories from the CME, which is realized via the SSA and Monte Carlo integration of these trajectories for moment estimation.

Second, we use the MoM, which derives a set of differential equations for the first *k* moments from the CME. These equations are not closed for systems with reactions of order two or higher, since lower-order moments depend on higher-order moments in these cases. We use truncation closure (TC) to solve this problem, a closure method that neglects the influences of higher-order moments on the considered set of moment equations. Unsurprisingly, it is, therefore, one of the most simplistic approaches, and we refer the interested reader to more sophisticated closure techniques that have been proposed to break this infinite hierarchy (see e.g. (12–14)). As an example, consider MoM approximations for the expected value 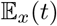 and variance 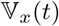 of the number of molecules *x* in a chemical system with only one species. After applying TC and thereby neglecting the influence of the skewness on the first two moments, the MoM equations read

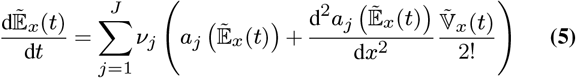

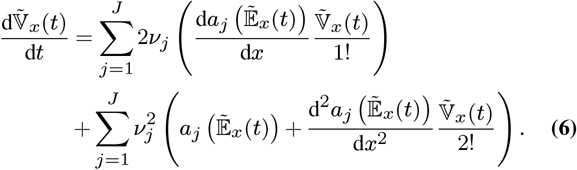

Here, the tilde denotes the Ordinary Differential Equation (ODE)-based approximation of the underlying quantity due to TC.

### Deriving a chemical reaction system that provably exhibits wildly stochastic behavior

In order to study the behavior of both moment estimation methods for systems that exhibit wildly stochastic behavior, we inversely design such a system. To this end, we consider the steady-state condition of the CME, which is obtained by setting the time derivative in Eq. (1) to zero and reads

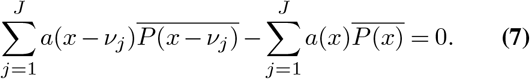

Reformulating our original goal, we would like to define a CRN whose equilibrium distribution 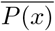 does not possess any statistical moments. One example of such a distribution on the natural numbers is given by

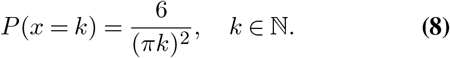

Equation Eq. (8) defines a proper, discrete, normalized probability measure on all natural numbers, as

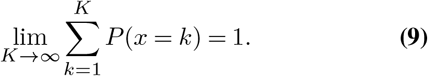

However, all moments of natural order *m* are defined by divergent sums,

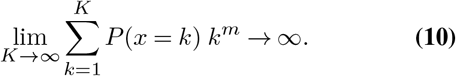

This holds regardless of whether one is interested in centralized moments or non-centralized counterparts.

The distribution defined in Equation Eq. (8) indeed describes the equilibrium distribution of a birth-death process with quadratic propensities,

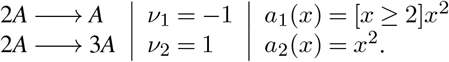

Here, [·] is a Boolean expression that is either equal to zero or one depending on the truth of the argument. Inserting our specific choice of system characteristics into Equation Eq. (7) results in

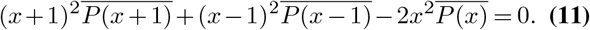

By replacing all three instances of 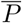 in Equation Eq. (11) using Equation Eq. (8), we obtain

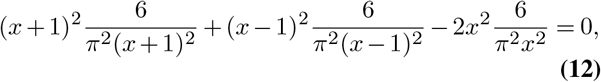

which holds for all *x*.

In summary, we defined a CRN whose equilibrium CME solution is a discrete heavy-tail distribution: It neither possesses an expected value nor any higher-order moment.

Our system is unphysical in two aspects. First, defining the propensities for second-order reactions according to the law of mass action, one counts the number of educt combinations, which is given by *x*(*x* – 1) instead of *x*^2^. This detail matters in particular for small molecule numbers. The second unphysical aspect is the Boolean expression that prohibits an absorbing system conformation at *x* = 0. In order to avoid this, we introduce a third reaction that slowly but constantly adds species *A* to the system. The resulting configuration transition graph of this modified system, which we refer to as System 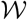, is shown in Figure 1A. It is more involved to derive the steady-state distribution for this altered system, which is why we resort to numerical simulations to validate our claims for realistic scenarios further. We nevertheless expect System 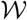 to have very similar characteristics because of nearly identical propensities, especially in the realm of many molecules. The time evolution of the system that is obtained from 25 000 SSA sample trajectories with initial configuration *x*(0) = 1 is illustrated in Figure 1B for five exemplary time points. As expected, a shift of the probability mass towards larger numbers of molecules can be observed.

**Fig. 1.**
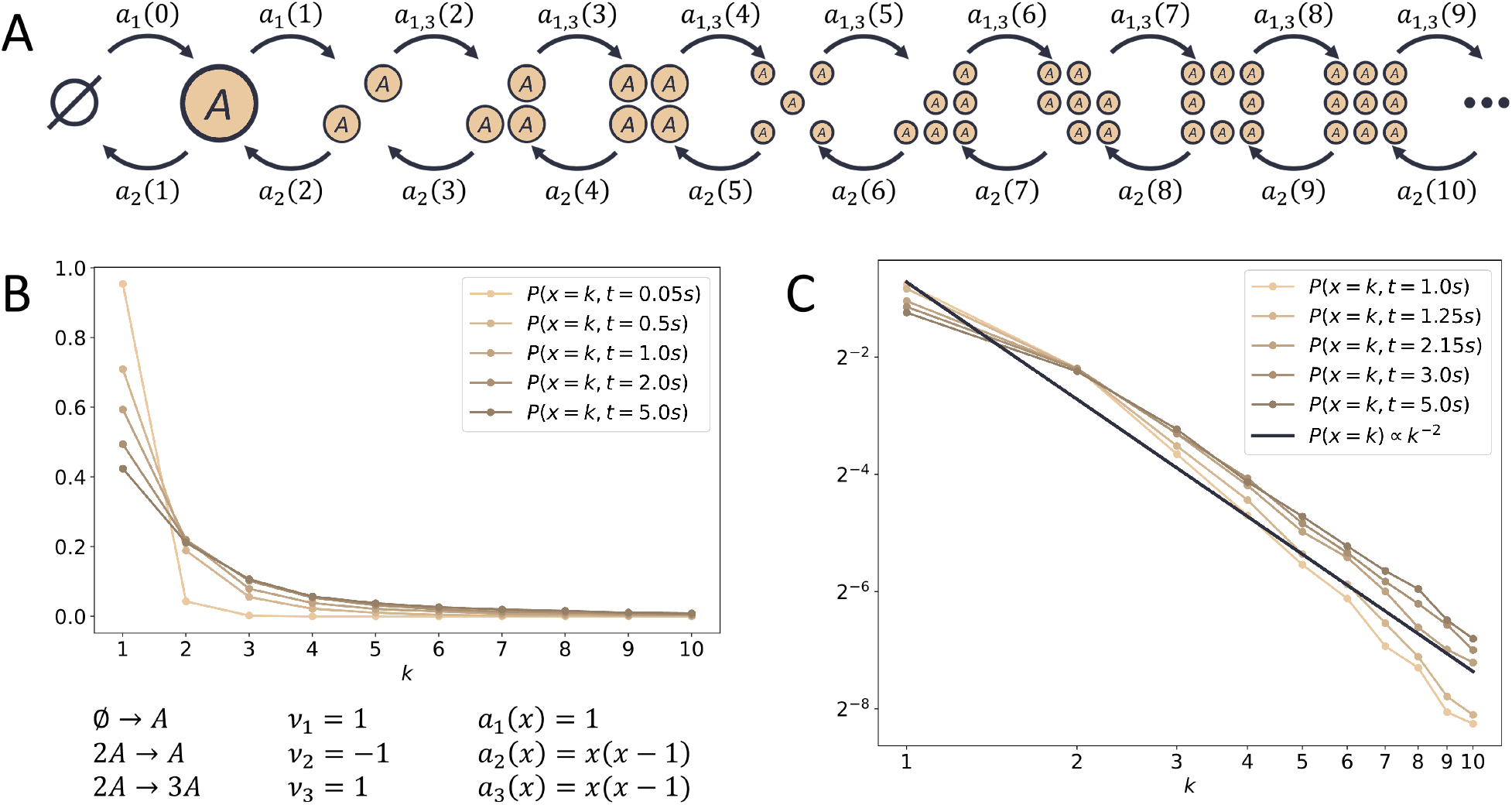
A birth-death process that exhibits wildly stochastic behavior. A. Configuration transition graph, B. *P*(*x,t*) estimated with 25 000 SSA trajectories that are evaluated over a logarithmically scaled, uniform time grid with 1 001 time points, *P*(*x,t* = 0) was set to *P*(*x* = 1, 0) = 1. C. A polynomial decay of distribution tails corresponds to a straight line in the double-logarithmic plot, as indicated with the black line of the heavy-tail distribution Eq. (8). The distribution *P*(*x,t*) also approaches such a straight line for later time points and *k* ≤ 2, thus indicating heavy-tailedness.

An obvious question at this point is whether System 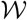 indeed possesses a heavy-tail steady-state solution. Figure 1C displays the evolution of the CME solution approximation of System 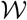 in comparison to the heavy-tail distribution Eq. (8). Since both axes are logarithmically scaled, the polynomial distribution Eq. (8) is a straight line with a defined slope. Distributions that decay faster than this reference correspond to lines with steeper slopes and vice versa. One clearly sees that the slopes of the CME solutions decay with growing time, eventually intersecting the slope of distribution Eq. (8). This point specifically marks the change from a CME solution with defined moments to a heavy-tailed one without. Therefore, the CME solution *P*(*x,t*) of System 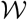 indeed makes a transition from an initial distribution with defined moments for early time instances to a heavy-tail distribution for later instances, and hence can be used for the analysis of moment-based approaches.

To put our findings into context, all simulations are similarly performed for a reference System 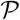 with well-defined moments depicted in Figure 2. This system is a simple immigration-death process with first-order kinetics for both reactions. The configuration transition graph can be adopted from System 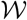 by neglecting the third reaction (which corresponds to setting *a_3_*(*x*) = 0). System 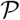 converges to a Poisson distribution with parameter *λ* = 10, i.e. the equilibrium distribution is 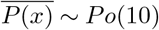, with expectation and variance 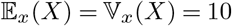.

**Fig. 2.**
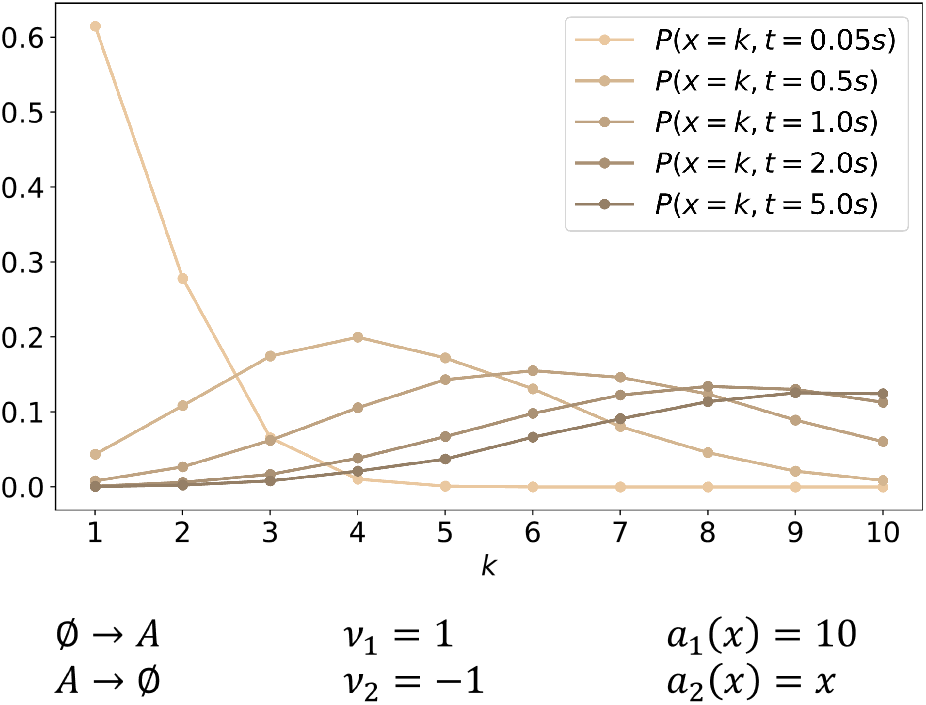
A simple immigration-death process with first-order kinetics serves as a benchmark problem. This system converges to a Po(10) distribution with well-defined moments. *P*(*x, t* = 0) has been set to *P*(*x* = 1,0) = 1.

### Moment estimators from SSA trajectories do not converge if the system exhibits wildly stochastic behavior

From the sampled trajectories of the CRN, one can of course still attempt to estimate the distributions’ statistical moments using the bias-free estimators

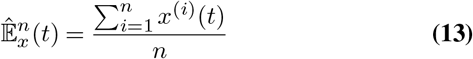

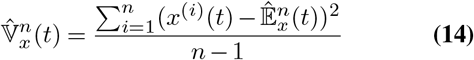

for the expected values 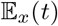 and variances 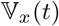. In this context, index *i* denotes one of the *n* independent SSA trajectories over which we average. Figure 3 compares these moment estimators for Systems 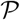 and 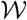 over an increasing number of sample trajectories for a time instance *t* = 5*s*. According to our analysis in Figure 1C, System 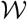 already exhibits wildly stochastic behavior at this point. Following the central limit theorem, one would expect 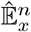 and 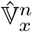 to converge for a growing number of included samples *n* in most scenarios. While this expectation is met for System 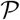 (Figure 3A), the same estimators applied to System 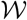 exhibit strikingly different behavior (Figure 3B): Since subsequent dots differ in exactly one additional sample taken into account during moment estimation, large gaps between subsequent points indicate disruptive trajectories that significantly influence the estimation of the whole trajectory ensemble and thereby prevent convergence. The most disruptive trajectory *x*^(17 895)^ of System 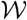 is visually highlighted and marks a huge gap for both moment estimators. Please note that the logarithms of both moment estimators have been visualized in order to cope with the large ranges these estimators cover. Summarizing, Figure 3 shows that statistical moment estimators do not converge for increasing numbers of SSA trajectories for System 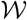 and cover a wide range of values.

**Fig. 3.**
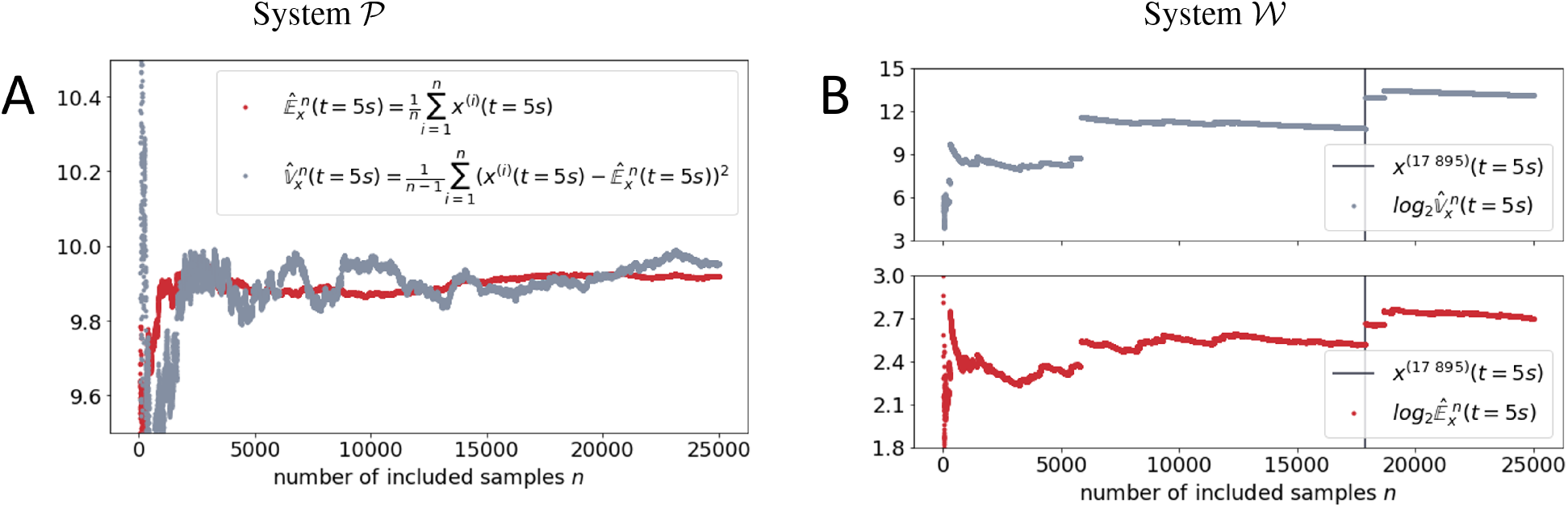
Moment estimators show qualitatively different behaviors for Systems 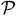 and 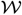 for a growing number of included sample trajectories. Bias-free moment estimators are applied to the final configuration of all sample trajectories *x*(*t* = 5*s*) for a growing number of included samples. Red and gray dots correspond to expectation and variance estimations, respectively. A. Results for System 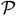 on a linear scale. B. Results for System 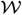 on a logarithmic scale. Trajectory 17 895 takes the value 10 554 at *t* = 5*s*.

Therefore, the estimation of moments from a fixed number of trajectories does not lead to reliable estimates.

### Expectation-based methods in heavy-tail applications

As indicated earlier, we are interested to test how the MoM behaves when applied to the two chemical Systems 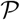 and 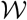. Using Equation Eq. (5) and Eq. (6), we derive the MoM ODEs for both simulated systems. For System 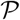, this leads to

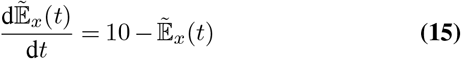

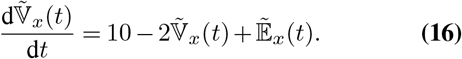

Similarly, the MoM ODEs for System 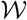 are derived as

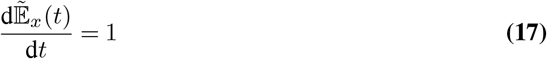

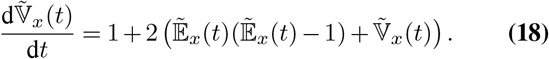

The just-presented equations are complemented with the deterministic initial conditions 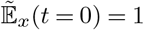 and 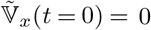, which correspond to the moments of the initial distribution *P*(*x* = 1, *t* = 0) = 1, to create solvable initial value problems. Figure 4 compares the time-dependent solution of the aforementioned equations to the bias-free moment estimators Eq. (13) and Eq. (14). Both the MoM solution and the sample-based estimators are evaluated over a grid of time points that are logarithmically equidistantly spaced so that time points are more densely distributed for smaller times. For the reference System 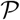 depicted in Figure 4A, the MoM prediction perfectly agrees with the sample-based estimators for the entire simulation time. Hence it can be assumed that both estimation techniques approximate the system’s characteristics properly.

**Fig. 4.**
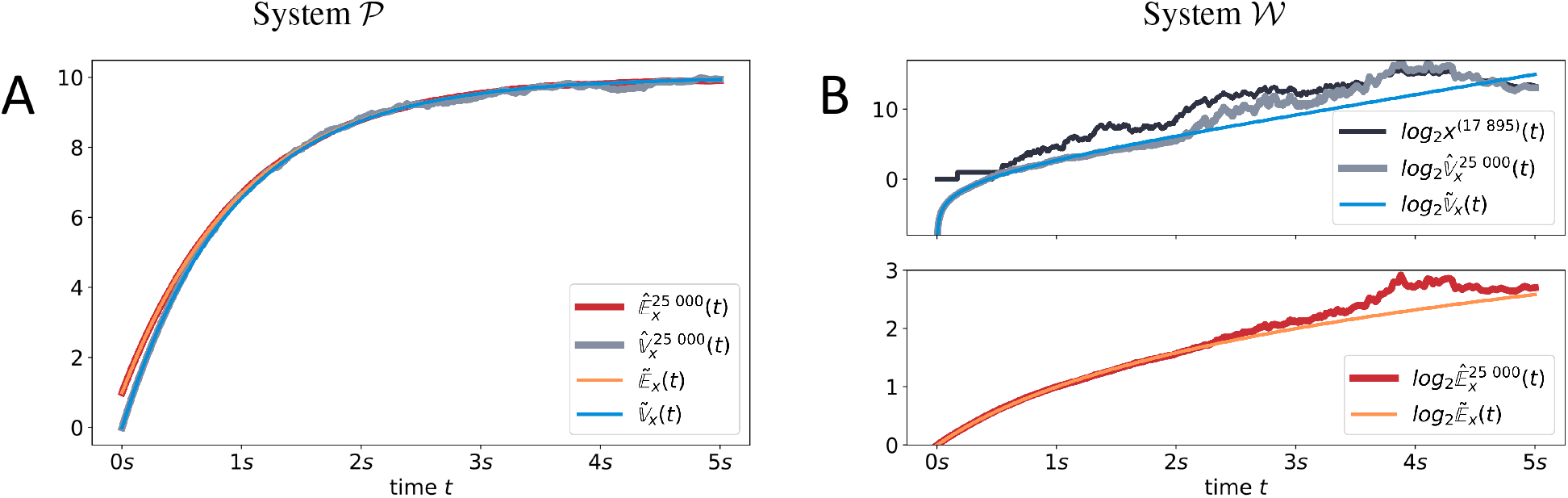
MoM results are qualitatively similar for both models, while sample-based estimators do not agree with them for System 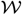, thereby indicating potentially non-existing moments. Comparison of time-dependent moment estimation for both simulated models. On the one hand, sample-based estimators for 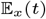 and 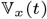 are depicted in red and light gray, respectively. MoM predictions, on the other hand, are visualized in orange and blue. As for Figure 3, trajectory *x*^(17 895)^(*t*) is plotted in dark gray to show how closely the sample-based variance estimation 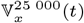 is related to this trajectory.

A qualitatively very different picture is painted for System 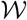 (Figure 4B). Initially, the two estimation techniques agree for both moments. However, after approximately one second, the sample-based estimation visually differs from the smooth MoM solution, especially for the variance. It is remarkable, that, although the moments still exist at that point in time, they are obviously increasingly difficult to estimate statistically. The quality of the sample-based estimation continues to decrease beyond the point, where System 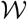 no longer possesses statistical moments.

The second observation in Figure 4 is less obvious at first but similarly essential: Even while crossing the point where the system’s statistical moments are no longer defined, the MoM trajectory is unaffected in nature. In other words, the MoM is unable to indicate the unpredictability of the underlying process.

Similar to Figure 3, we plotted the time course of sample trajectory *x*^(17 895)^(*t*). Besides the order of magnitude of the number of simulated molecules, it is remarkable how similar the variance estimation 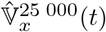 resembles this trajectory, especially after 2 seconds of simulated time. Hence, this single sample dominates the overall estimation process, which inherently questions the usability of statistical moments in such a context.

In summary, results show that the application of the MoM to CRNs that show wildly stochastic behavior provides smooth moment estimates that are similar to moment courses of systems with well-defined moments. Hence, MoM moment trajectories are not indicative of heavy-tailedness. In contrast, the consistency of moment estimates from Gillespie trajectories is lost over time, and single Gillespie trajectories with large molecule numbers dominate the estimates of both moments. A visual indicator of this is the increasing ruggedness of the estimated moment trajectories over time.

### Wildly stochastic behavior results in heavy-tailedness of simulation run-times

Our results outlined the difficulties of identifying and predicting wildly stochastic CRNs. The two challenges clearly are related, as the identification of heavy-tailed CME solutions is the basis for a subsequent adequate choice of approximation methods. In this chapter, we would like to present a diagnosis approach while simultaneously discussing another interesting aspect of wildly stochastic behavior in the context of CRNs.

To this end, we revisit the SSA used to draw sample trajectories from the CME solution. Starting from the initial configuration *P*(*x,t* = 0), the algorithm samples the index *j* of the next reaction and a waiting time *τ*. The waiting time is drawn from an exponential distribution 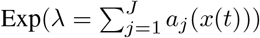. This distribution has expectation value *λ*^-1^.

The number of needed SSA steps to reach 5 seconds of simulated time is displayed as a histogram over all 25 000 sampled SSA trajectories for both systems in Figure 5. For System 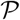 (Figure 5A), the histogram resembles a discretized Gaussian distribution with a mean of approximately 90 SSA steps per simulation. Moreover, the probability of one simulation trajectory with less than 40 or more than 140 steps within 5*s* of simulated time is approximately zero. In contrast to this, the histogram depicted in Figure 5B is remarkably different in several aspects. Firstly, it is overall more likely that System 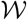 needs an odd number of steps to surpass 5*s* of simulated time. While this may sound surprising at first, it can be related to the very small sum of propensities associated with configuration *x* = 1 that translates to a long resting time. In fact, Figure 1B shows that over 40%of all SSA trajectories finish in this configuration. No matter how their journey through configuration space went until then, they are bound to need an odd number of reactions to surpass the point of 5*s*. The second aspect worth discussing is the long histogram tail in Figure 5B. It originates from the two quadratic propensities that lead to vanishing expected time increments for large numbers of simulated molecules. Different realizations of System 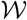, therefore, exhibit completely different behaviors: While some trajectories only take one SSA step to reach 5 seconds of simulated time, other trajectories escape the regime of small molecule numbers (and therefore propensities). Once a certain number of molecules is reached, the system performs an incredible number of reactions to progress only slightly in time. The probability of such an escape from small molecule numbers can be deduced from the black cumulative distribution curve in Figure 5B. It indicates that more than 20%of all SSA samples are not finished after 175 reactions.

**Fig. 5.**
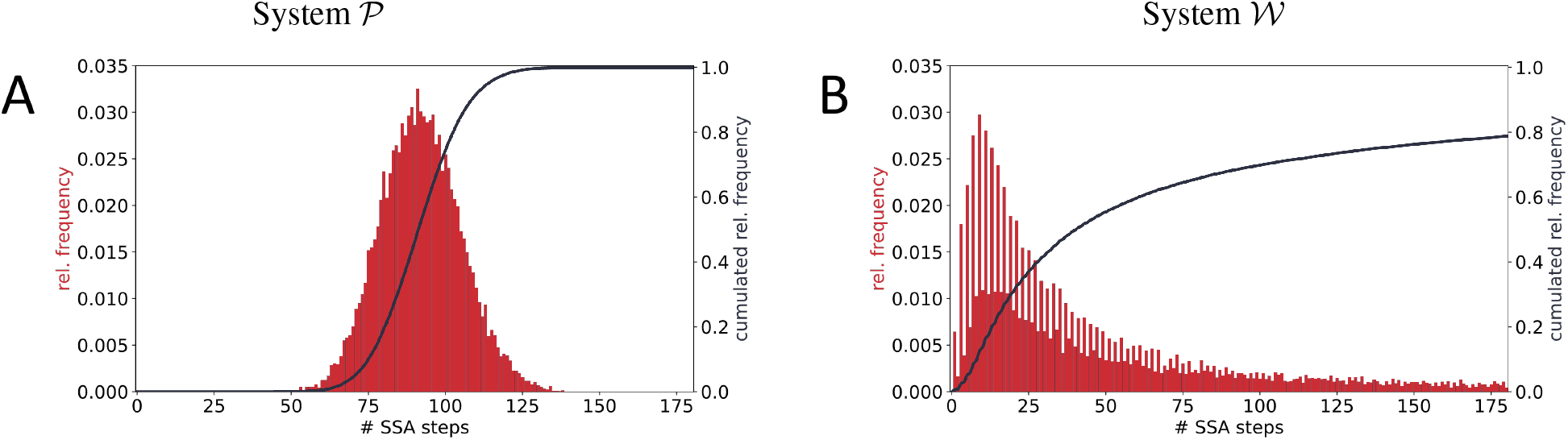
Heavy-tailedness of the CME solution translates to the distribution of run times. The histograms describe the number of reactions needed to simulate 5*s* of simulated time. Both systems are simulated 25 000 times. The relative frequency is depicted as a red histogram while its cumulative distribution is visualized as a black line. The two quantities are plotted on different scales. A. Results for Systems 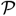, B. Results for System 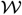.

The number of performed SSA steps is directly correlated with the simulation run-time. A scientist attempting to draw samples from a heavy-tailed CME solution will therefore experience many very fast simulations at first. However, the overall run-time is completely dominated by single extreme events. The simulation of trajectory *x*^(17 895)^(*t*), for instance, took several days, while most of the 25 000 samples were completed within a fraction of a second. Besides the unpleasant observation of such extreme samples, the distribution of run-times like the one depicted in Figure 5 can already unveil the heavy tails of the CME solution itself.

## Discussion and Conclusion

The CME is widely recognized for accurately describing the behavior of CRNs of various different types and characteristics. Due to this flexibility, it is in principle not surprising that the CME solution of particular systems does not possess any statistical moments. However, it still is counter-intuitive that those systems can be derived from simple mass action kinetics of second-order reactions.

In this work, we first derived a CRN that provably converges to a heavy-tail distribution. After slightly altering the system to increase interpretability, we numerically validated our theoretical claims. On this basis, we presented evidence that expectation-based CME approximation methods are prone to fail when applied to systems that converge to heavy-tail distributions. Moreover, the MoM does not indicate its failure by any means. In our eyes, it is, therefore, crucial to examine each CRN thoroughly before choosing an adequate approximation technique for the CME. To this end, we propose to employ a sample of SSA simulations. The corresponding implementation is simple, yet, SSA trajectories represent a true sample of the underlying CME and are therefore capable of predicting different actual system trajectories. Applying moment estimators to the sampled trajectories and observing their convergence can furthermore indicate whether or not statistical moments of the CME solution exist and how easily they can be estimated. Finally, one interesting aspect of the SSA is that its run-time scales linearly with the number of recorded reactions. In the context of our simulations, extreme events that effectively prevent the CME solution from having statistical moments correspond to simulations where the system escapes the slow dynamics of a few molecules, thereby further increasing the reaction propensities quadratically. As indicated earlier, this vast amount of reactions manifests itself through surprisingly large simulation run-times. Effectively, the heavy tail of the CME solution is therefore transferred to the distribution of simulation run-times, which is a quantity most simulation researchers attentively monitor and which therefore serves as another criterion to assess the system’s expectability.

## Abbreviations

(CME): Chemical Master Equation
(CRN): (Bio-)Chemical Reaction Network
(MoM): Method of Moments
(ODE): Ordinary Dif-ferential Equation
(SSA): Stochastic Simulation Algorithm
(TC): truncation closure

## Availability of data and materials

All code is available via FAIRDOM Hub and can be executed with any common *Python 3* distribution. Necessary packages are publicly available.

## Authors’ contributions

Conceptualization, V.W and N.R.; Methodology, V.W.; Software, V.W.; Validation, V.W. and N.R.; Formal Analysis, V.W. and N.R.; Investigation, V.W. and N.R.; Resources, N.R.; Data Curation, V.W.; Writing—Original Draft Preparation, V.W.; Writing—Review and Editing, V.W. and N.R.; Visualization, V.W. and N.R.; Supervision, N.R.; Project Administration, N.R.; Funding Acquisition, N.R.

## Acknowledgments

This preprint was formatted using a LATEX class by Ricardo Henriques that can be accessed here.

